# Horizontal gene transfer by plasmids relaxes the evolutionary constraints of its own tradeoff

**DOI:** 10.64898/2026.05.31.729049

**Authors:** Olivia Kosterlitz, Elizabeth S. Duan, Maya Abhyankar, Benjamin Kerr, Eva M. Top

## Abstract

Horizontal gene transfer (HGT) is a defining feature of plasmid biology, enabling plasmids to spread between bacterial cells and driving the global dissemination of antibiotic resistance and other adaptive traits. HGT and vertical gene transfer (VGT) have long been assumed to be subject to an evolutionary tradeoff, where improvements in one come at the expense of the other. Yet whether this tradeoff reliably constrains plasmid evolution remains unclear. Through the first cross-literature synthesis examining both transmission traits across 16 studies and 245 plasmid-host pairs, we find mixed empirical evidence: patterns consistent with a tradeoff alongside outcomes that a strict tradeoff should make impossible. We propose that this contradiction is resolved by recognizing that a tradeoff is ensured only when plasmid-host pairs are well-adapted to one another. Because HGT introduces plasmids into novel hosts where this adaptation is disrupted, it systematically creates the very conditions under which near-term evolution need not be bound by a tradeoff. We support this framework empirically by documenting the first mutation that simultaneously improves both transmission modes, arising from a non-coevolved plasmid-host pair. These findings reveal a fundamental irony: the defining feature of plasmid transmission is the very mechanism that relaxes the evolutionary constraints of its own tradeoff.

## Introduction

The rapid spread of adaptive traits such as antibiotic resistance across bacterial communities is driven largely by plasmids^1,2^. These mobile genetic elements spread via two modes of transmission: vertical gene transfer (VGT), in which plasmids are inherited from mother to daughter cell following binary fission, and horizontal gene transfer (HGT), in which plasmids are transferred between different cells including distantly related species. Because both modes generate new plasmid copies, VGT and HGT together constitute the two core components of plasmid fitness. It follows that if both the VGT and HGT rates were free to evolve independently, selection would favor increases in both until each reached its maximal value in a given host. However, HGT by plasmid conjugation can be costly^3,4^, reducing resources available for host growth and lowering VGT relative to a plasmid-free cell. This has motivated the hypothesis that VGT and HGT are subject to an evolutionary tradeoff^5–8^, where improvement in one transmission mode cannot occur without a decrease in the other, a widely invoked constraint on plasmid evolution^9–11^.

The empirical record of a VGT-HGT tradeoff is mixed. Evidence consistent with the tradeoff exists: a negative correlation between transfer rate and host fitness is observed across diverse plasmid-host pairs^9^, and evolution experiments have produced antagonistic outcomes in which one transmission mode improves at the expense of the other^5,12^. Yet, contradictory evidence also exists. Compensatory mutations that ameliorate plasmid cost can increase VGT through pathways entirely unrelated to and not affecting HGT^13–15^, and mutations that increase HGT have been documented with no detectable effect on host fitness^16^. These observations indicate that both traits can improve independently of one another, an outcome inconsistent with a tradeoff. This contradiction has been noted but not explained^17,18^, leaving the most consequential question unresolved: under what conditions, if any, does a VGT-HGT tradeoff apply?

This question has consequences beyond basic biology. Evolutionary tradeoffs inform evolutionary predictions and are therefore critical to formulating strategies in evolutionary medicine and ecology to steer evolutionary outcomes toward desired ends. The VGT-HGT tradeoff in particular has been invoked as relevant to limiting the spread of antibiotic resistance plasmids^5,9,19,20^. If improving one transmission mode necessarily harms the other, evolutionary options are restricted and interventions become more predictable. Yet the predictive value of any tradeoff depends entirely on whether and when it constrains evolutionary outcomes, a condition that has not been rigorously evaluated for the VGT-HGT system.

We propose that the key to resolving the contradictory evidence for the VGT-HGT tradeoff lies in how well a plasmid is adapted to its host. Consider a plasmid recently introduced into a novel host: such a pair is likely suboptimal for both transmission modes, with room to improve in both simultaneously. By contrast, a plasmid long coevolved with its host will have likely exhausted those opportunities, such that any improvement in one trait necessarily comes at the expense of the other. The degree to which plasmid evolution is constrained to such antagonistic outcomes, therefore, depends on the starting performance of the ancestral plasmid-host pair. This reasoning leads to an interesting twist: HGT itself may be a primary mechanism that relaxes its own tradeoff by continuously generating suboptimal plasmid-host pairs through conjugation-mediated introduction of the plasmid into non-coevolved hosts. We explore these ideas through the first cross-literature synthesis to examine both transmission traits across 16 studies, and through new phenotypic data from our previously published evolution experiment^21^. Together, these analyses suggest that evolutionary change consistent with a VGT-HGT tradeoff may be systematically relaxed by HGT itself. Most directly, we document the first mutation that simultaneously improves both transmission modes, a category of outcome inconsistent with a classical tradeoff but consistent with a poorly adapted plasmid-host pair. These findings reshape our understanding of how plasmid transmission evolves and have direct implications for strategies that rely on the VGT-HGT tradeoff to predict and limit the spread of antibiotic resistance plasmids.

## Results

### A trait space framework clarifying when tradeoff predictions apply

We represent VGT and HGT as a two-dimensional trait space, with HGT on the horizontal axis and VGT on the vertical axis (Figure 1). Both traits directly contribute to plasmid reproduction such that higher values along either axis increase overall plasmid fitness. If both traits were unconstrained, all combinations of VGT and HGT would be possible. However, under the logic of costly HGT, some trait combinations are impossible (grey region across panels, Figure 1). No genotype can simultaneously maximize both traits (joint maximum, Figure 1a). We assume the interface between possible and impossible regions of trait space is a continuous decreasing function, which we hereafter refer to as the “boundary” (concave curve across panels, Figure 1). These assumptions about the structure of trait space are shared with a few recent frameworks across biological systems^22–24^, which we apply here to the VGT-HGT system.

**Figure 1.**
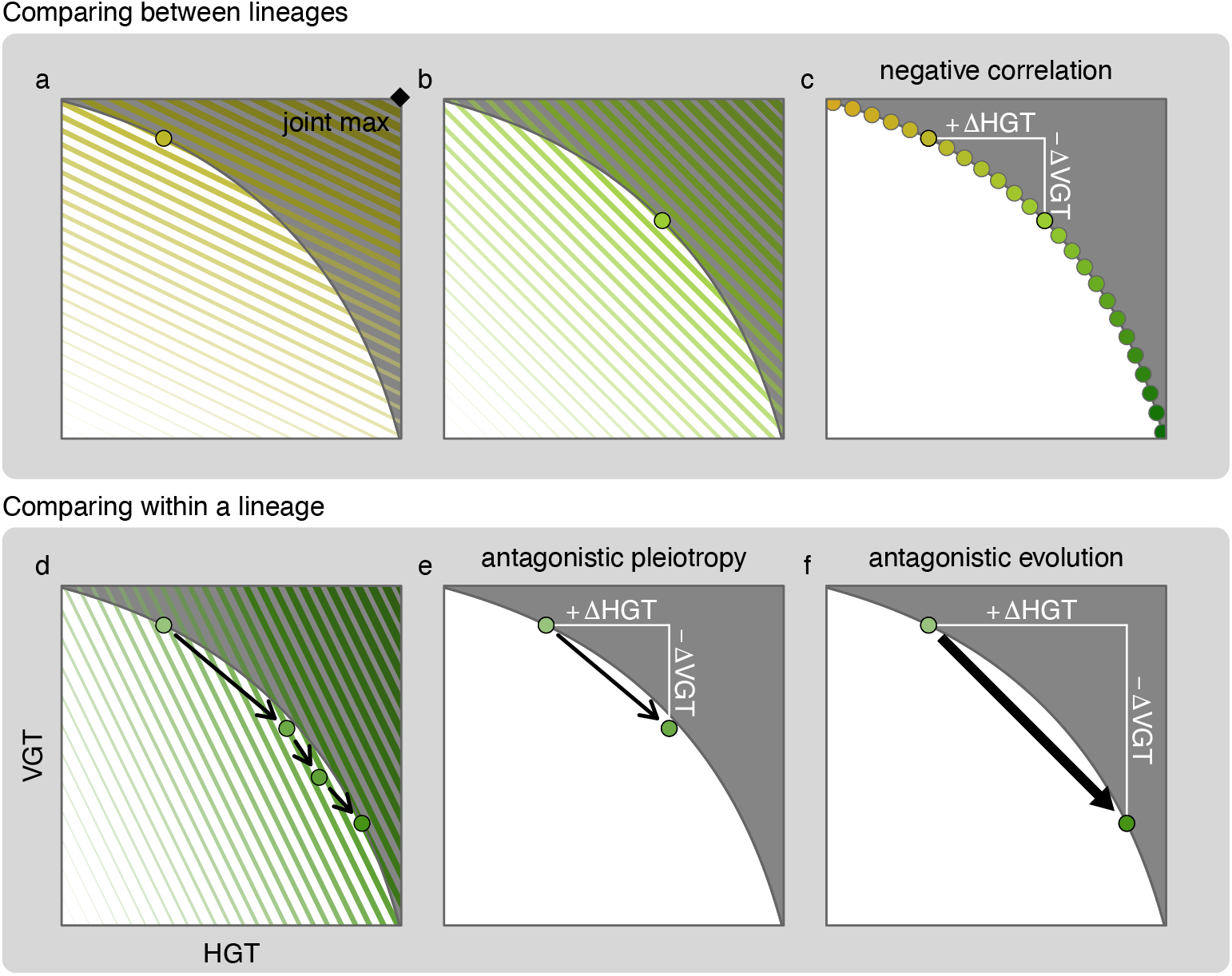
Schematic demonstrating the three common empirical signatures of an evolutionary tradeoff. All panels share the same trait space structure: HGT rate on the horizontal axis, VGT on the vertical axis, a boundary (negatively sloped curve) separating the impossible trait space (dark grey) from the possible trait space (white). (a) The joint maximum is marked by a diamond. (a-b) Two environments favoring different trait combinations direct evolution toward different optima on the boundary, each corresponding to the point of tangency between the colored fitness contours and the boundary. Colored lines show fitness variation across trait space, with darker and thicker lines indicating higher fitness. (c) Across many lineages evolving under different environments, genotypes accumulate along the tradeoff boundary, producing a negative correlation between traits, a common empirical signature of an evolutionary tradeoff. (d) A lineage evolving within a single environment acquires successive mutations (black arrows) that move it toward the fitness optimum on the boundary. (e) At the shortest timescale, comparing an ancestor to the first beneficial mutation reveals antagonistic pleiotropy: one trait increases at the expense of the other (e.g., +ΔHGT, −ΔVGT). (f) At the longest timescale, comparing an ancestor to a multi-mutation descendant reveals antagonistic evolution, the same antagonistic signature accumulated across multiple mutations.

In this trait space, fitness is maximized at the boundary. For any given environment, the fitness optimum corresponds to the point where the selection gradient is tangent to the boundary (Figures 1a and b). Natural selection therefore drives lineages toward the boundary, such that well-adapted lineages are expected to reside on or near it. Comparing two lineages on the boundary produces a negative correlation (Figure 1c), one of the most widely used empirical signatures consistent with a tradeoff^25^.

The same negative relationship arises when following a lineage across time. If the environment changes, a previously well-adapted genotype will evolve toward the new fitness optimum, which lies at a different point on the boundary (Figure 1d). This evolutionary trajectory produces two additional empirical signatures of a tradeoff^26^, depending on the timescale considered. At the shortest timescale, comparing the ancestor to the first beneficial mutation reveals antagonistic pleiotropy (Figure 1e), a mutation that increases one trait at the expense of the other. At a longer timescale, comparing the ancestor to a descendant carrying multiple mutations reveals antagonistic evolution (Figure 1f).

These three common empirical signatures share a fundamental feature: each evaluates the tradeoff as a pairwise comparison between two points in trait space, where one trait value has increased and the other decreased. This applies whether the comparison is across lineages at a single time point (Figure 1c) or within a lineage across different points in time (Figures 1e and f). In all the cases considered so far, at least one point of comparison lies on the boundary. Importantly, when this boundary residency holds, these empirical signatures of a tradeoff are guaranteed.

This framework explains why mixed empirical signatures of a tradeoff can arise: when boundary residency does not hold, these empirical signatures need not follow. Indeed, the existence of any one of the three signatures (Figures 1c, e, and f) does not guarantee the other two will be found. To illustrate, we focus on within-lineage comparisons across time, the comparison commonly made in laboratory evolution experiments. Evolutionary outcomes are categorized using a Cartesian coordinate system with the ancestor at the origin and trait changes in the descendant on each axis (Figure 2a). Three outcome categories are possible. Antagonistic outcomes occur when one trait improves while the other decreases (Q2 and Q4, Figure 2a); for a single mutation this is antagonistic pleiotropy, and across multiple mutations this is antagonistic evolution. Synergistic outcomes occur when both traits improve simultaneously (Q1, Figure 2a). Single-trait outcomes occur when one trait changes with no detectable effect on the other (A1 and A2, Figure 2a); for a single mutation, these are non-pleiotropic mutations. The latter two categories are inconsistent with a tradeoff, but, importantly, they remain possible within the structural assumptions of trait space.

**Figure 2.**
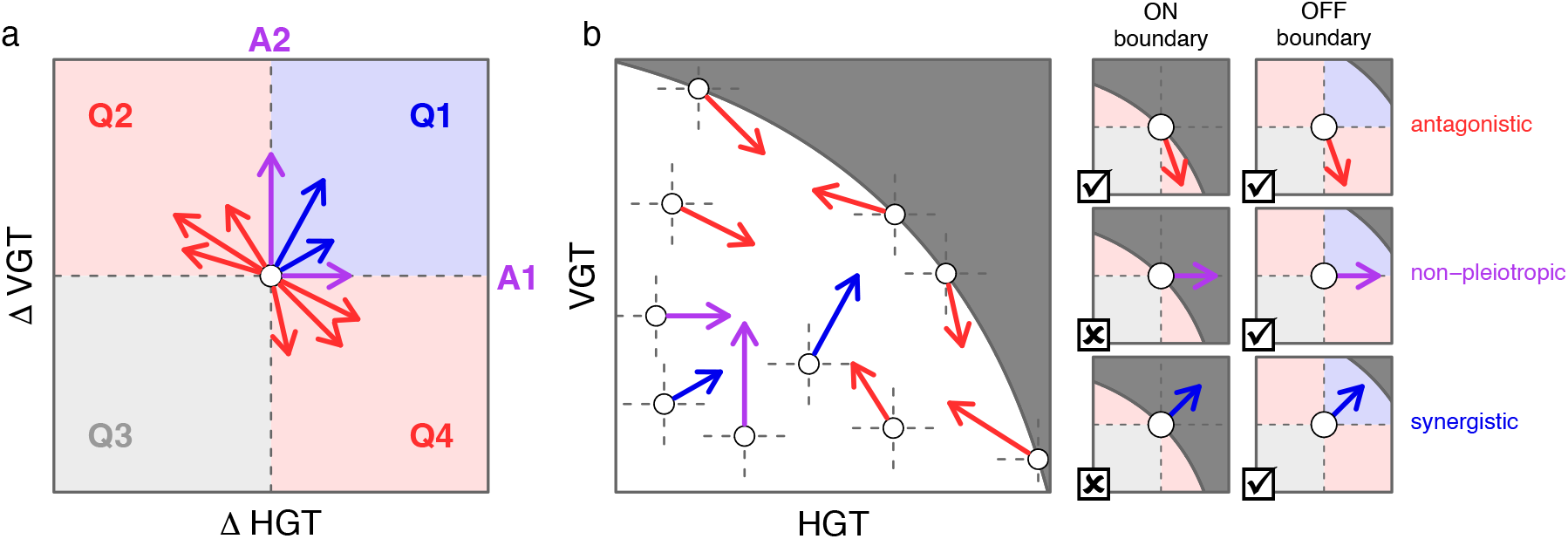
Relative evolutionary outcomes (a) mapped to absolute trait space (b) reveal that starting position in trait space determines mutational accessibility. (a) Possible evolutionary outcomes represented as a Cartesian coordinate system with the ancestor at the origin and changes in HGT and VGT on the horizontal and vertical axes, respectively. Outcomes are classified by quadrant: synergistic (Q1, blue), antagonistic (Q2 and Q4, red), and both-traits-decreasing (Q3, light grey). Since Q3 decreases both traits and is not selectively favored, we omit this category of outcome throughout the manuscript. Single-trait outcomes fall on the axes (A1, A2, purple). (b) Mapping the relative outcomes given in panel a into absolute trait space using ancestral trait values reveals that accessible evolutionary outcomes depend strongly on ancestral position. Genotypes on the boundary have access only to antagonistic pleiotropy (red arrows), while interior genotypes retain access to synergistic (blue arrows), and non-pleiotropic (purple arrows) mutations. The dark grey region denotes impossible trait combinations. Small panels on the right illustrate outcome accessibility for genotypes on versus off the boundary: a checkmark indicates the mutation lands in possible trait space and is therefore accessible, while an X indicates the mutation would require entering impossible trait space and is therefore inaccessible.

This is well illustrated by placing these relative comparisons into trait space using the absolute trait values of the ancestor (Figure 2b). Genotypes on the boundary have access only to antagonistic mutations — every accessible mutation that improves one trait must decrease the other — and are therefore subject to a strict tradeoff. Interior genotypes, by contrast, retain access to synergistic and non-pleiotropic mutations, and a tradeoff is not guaranteed. The implication is that a tradeoff is conditional and may depend on a lineage’s position in trait space, an insight shared with similar frameworks in other biological systems^23,24^.

### Boundary departure disrupts tradeoff predictability

Genotypes off the boundary are therefore consequential: their evolutionary trajectory is not constrained to antagonism alone. This raises an obvious question: what processes displace lineages into the interior? Existing frameworks have focused on selection moving lineages toward the boundary, with less attention on how departure occurs. Yet because off-boundary genotypes can evolve with fewer constraints, establishing how often lineages visit the interior of trait space is important.

Somewhat counterintuitively, departure from the boundary can occur through beneficial mutations. It is intuitive that mutations move lineages closer to the boundary, since the boundary contains the highest fitness genotypes across environments (Figure 1c). However, in any single environment, only a subset of boundary positions are high fitness (Figure 3a). Therefore, a beneficial mutation need not land on or near the boundary, and may fall well within the interior (Figure 3b). Once off the boundary, synergistic and non-pleiotropic mutations become accessible, and subsequent evolution is no longer constrained to antagonism alone (right panel, Figure 3b).

**Figure 3.**
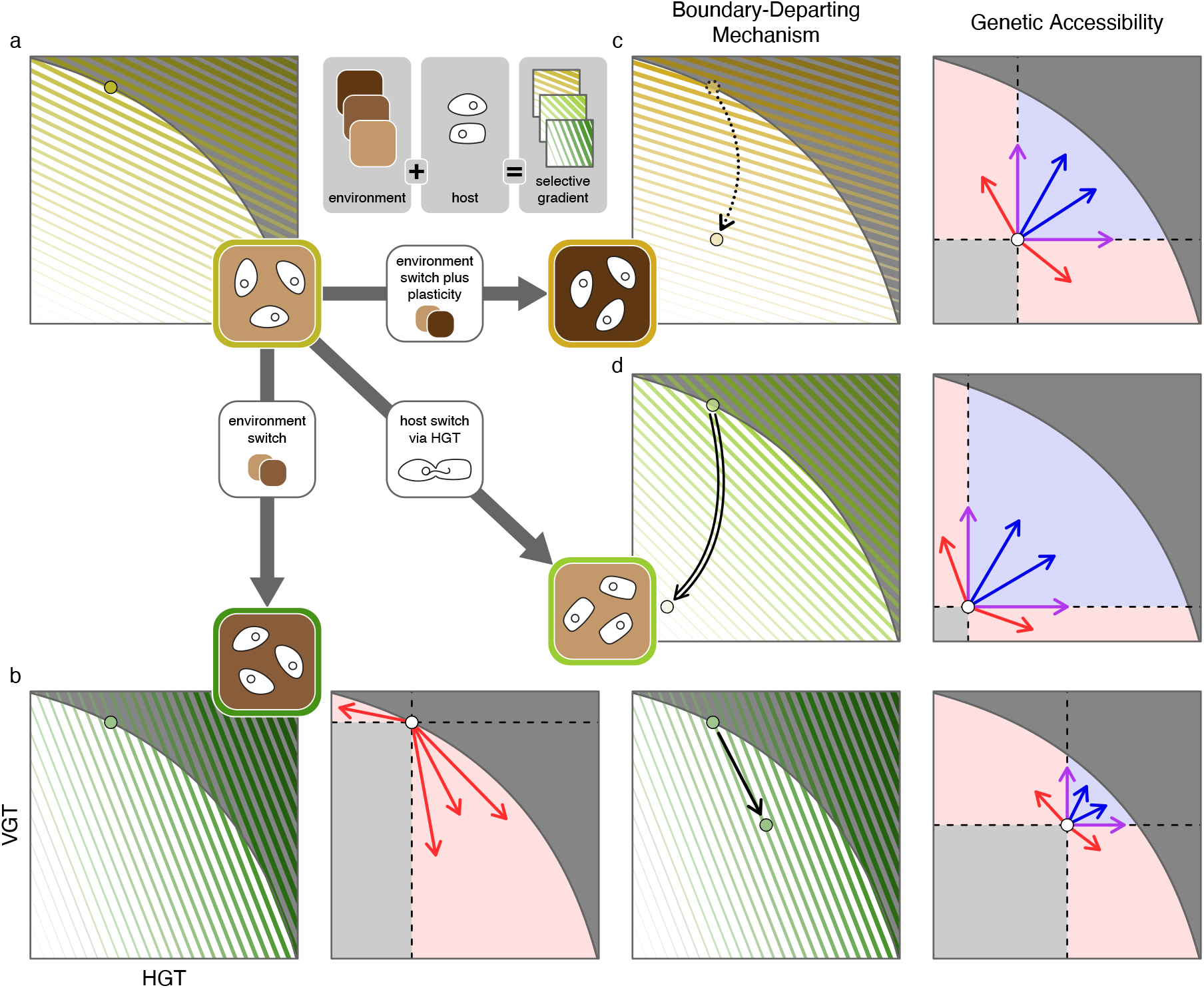
General mechanisms of boundary departure and their evolutionary consequences. The schematic key (top center) illustrates that the selection gradient for the plasmid in trait space is determined jointly by the environment (shades of brown) and host background (cell shapes). (a) A plasmid-host pair resides at a fitness optimum on the boundary under an initial environment. (b) An environmental shift alters the selection gradient (from light yellow lines in panel a to dark green lines in panel b), moving the fitness optimum to a new location on the boundary (left subpanel). Because the lineage initially resides on the boundary, all accessible beneficial mutations exhibit antagonistic pleiotropy (second subpanel). Fixation of one such mutation increases fitness but displaces the lineage into the interior (third subpanel), expanding genetic accessibility to include synergistic and non-pleiotropic mutations (fourth subpanel). (c) Phenotypic plasticity accompanying an environmental shift causes immediate displacement off the boundary (dotted arrow) before any mutation occurs, similarly expanding genetic accessibility. (d) HGT introduces the plasmid into a novel host background, creating a new lineage off the boundary (double-edge arrow), again expanding genetic accessibility. Right-hand panels in (b), (c), and (d) illustrate accessible mutations from the post-displacement interior position. All panels are schematics intended to illustrate general features of each mechanism.

Boundary departure can also occur through phenotypic plasticity, where the same genotype expresses different trait values in different environments (Figure 3c). A shift in the environment can therefore displace a genotype from the boundary before any mutation occurs, and the first beneficial mutation need not exhibit antagonistic pleiotropy.

It is noteworthy that the VGT-HGT tradeoff has a unique boundary-departing mechanism: horizontal gene transfer itself (Figure 3d). HGT events place plasmids into novel host backgrounds, disrupting long-term coevolutionary adaptation and violating the assumption that lineages have had sufficient time under stable selection to reach their fitness optimum. Empirically, introducing plasmids into novel hosts increases plasmid cost, reducing VGT^3^. Subsequent compensatory evolution in the plasmid, the host, or both acts to reduce this cost^15^. Together, these observations imply that HGT systematically creates plasmid-host pairs in the interior of possible trait space, away from the boundary, relaxing evolutionary constraints.

Because HGT is ubiquitous, plasmid-host pairs are regularly created off the boundary before compensatory evolution can act. Boundary residency may therefore be transient or absent for many plasmid-host pairs. This matters because the classical tradeoff expectation of antagonistic evolution follows from boundary residency. Regular creation of off-boundary pairs by HGT means these predictions can frequently fail in practice. This is not a rejection of a VGT-HGT tradeoff itself, but a challenge to the assumption that lineages reside on the boundary consistently enough to generate reliable tradeoff predictions.

### Negative correlation across plasmid-host pairs is consistent with a VGT-HGT tradeoff

We next asked whether empirical measurements of VGT and HGT across diverse plasmid-host pairs are consistent with the trait space framework. We compiled 16 published studies measuring both traits, comprising 245 plasmid-host pairs spanning more than 14 replicon types, 3 MOB groups, and 5 host species (Extended Data Figure 1, Extended Data Figure 2 & Supplementary Table 1)^4,5,9,12,13,16,18,27–35^. Of these, 196 pairs report absolute measurements of both transfer rate and host fitness and can be plotted directly in trait space (Figure 4a). The remaining pairs report only relative changes between ancestors and descendants from evolution experiments and are analyzed in the following section. HGT values, measured as plasmid transfer rates, span ten orders of magnitude (10^−18^ to 10^−8^). VGT values, measured as host fitness relative to a plasmid-free host, range from 0.3 to 1.5, reflecting plasmids that range from imposing substantial fitness costs to conferring large fitness benefits.

**Figure 4.**
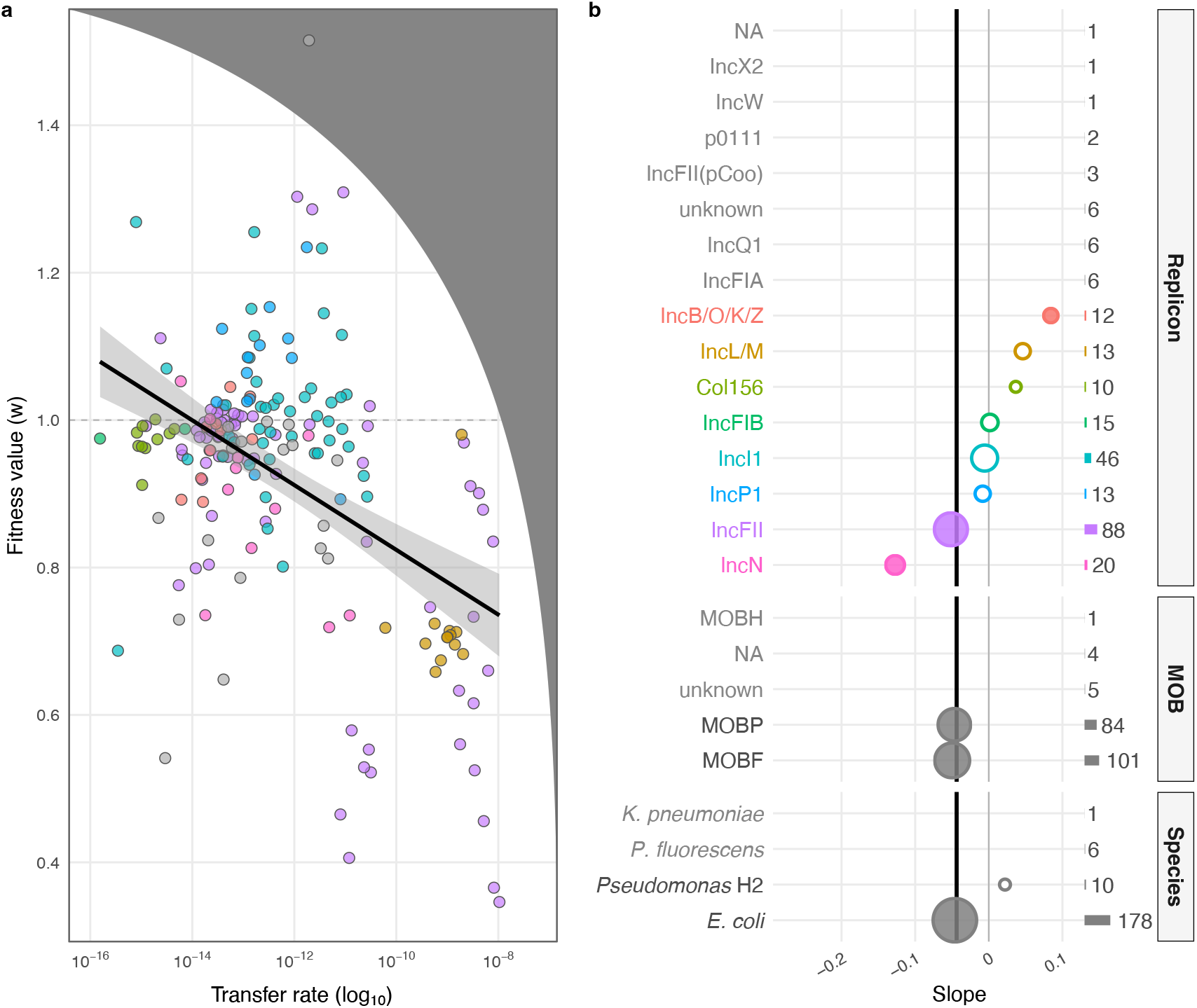
Negative correlation between vertical and horizontal transmission across diverse plasmid-host pairs. (a) Cross-literature comparison of plasmid-host pairs (n = 196) measuring HGT given as plasmid transfer rate (x-axis, log10 scale) and VGT as plasmid-bearing host fitness relative to a plasmid-free host (y-axis). The dashed horizontal line at w = 1 denotes the plasmid-free host fitness; values above indicate that the plasmid confers a fitness benefit and values below indicate fitness costs. Points are colored by replicon type (see panel b); replicon groups with fewer than 10 members are shown in grey. The black regression line (p < 0.001) with 95% confidence interval (grey shading) indicates a significant negative correlation. The hypothetical schematic boundary and impossible trait region (dark grey, upper right) are overlaid to connect empirical observations to the trait space framework in Figures 1-3. (b) Slopes of the VGT-HGT relationship (x-axis) estimated separately for three categories: replicon type, MOB type, and host species. The vertical black line indicates the overall slope from panel a; the vertical grey solid line at zero indicates the null expectation of no correlation. Point size, horizontal bars, and labeled numbers indicate group sample size; subgroups with fewer than 10 members were excluded from analysis. Filled circles indicate statistically significant correlations. Nearly all significant subgroup correlations are negative, with the exception of the replicon group IncB/O/K/Z.

Plotting this subset in absolute trait space reveals several patterns (Figure 4a). Pairs do not occupy the joint high-transfer, high-fitness region, consistent with an impossible region of trait space (Figure 1). A significant negative correlation across this dataset is consistent with a negatively sloped boundary, an empirical signature of a tradeoff (Figure 1c). However, a majority of plasmid-host pairs lie well within the interior of the possible region rather than clustering near a boundary. This interior occupancy is consistent with HGT-driven boundary departure, where plasmids introduced into novel hosts have not yet undergone sufficient compensatory evolution to become well adapted (Figure 3d). Phenotypic plasticity from lab measurement conditions may also contribute, displacing plasmid-host pairs from their in situ trait values (Figure 3c).

At the subgroup level, patterns are more mixed (Figure 4b). Among replicon groups with more than 10 pairs, only 3 show significant correlations: IncFII and IncN show negative slopes while, interestingly, IncB/O/K/Z shows a positive slope. Both MOB groups with sufficient data, MOBF and MOBP, show negative slopes, as does *E. coli*, which comprises the large majority of pairs. Most subgroup estimates are imprecise given small sample sizes, precluding strong conclusions at this level.

### Evolution experiments produce non-antagonistic outcomes inconsistent with classical tradeoff predictions

Plasmid evolution experiments allow us to test whether evolutionary responses are consistent with classical tradeoff predictions of antagonistic evolution (Figure 5). We classify each ancestor-descendant pair into the three evolutionary outcome categories (antagonistic, synergistic, or single-trait effects; Figure 2a). If ancestors reside on a boundary, all evolutionary responses should be antagonistic (Q2 and Q4; Figure 2b). Outcomes where only one trait changes (A1, A2) or both traits improve (Q1) are only possible if ancestors start away from a boundary (Figure 2b).

**Figure 5.**
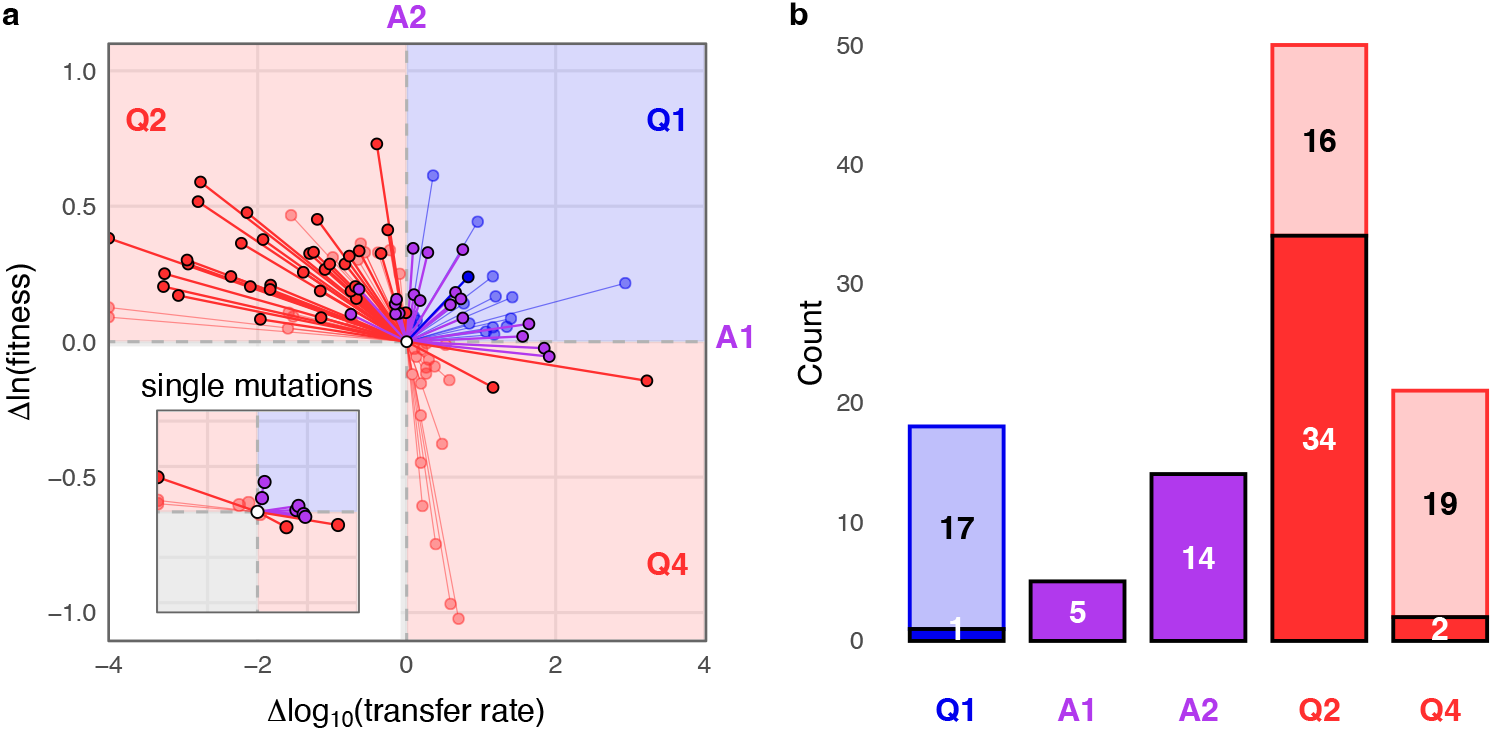
Evolutionary outcomes from plasmid evolution experiments reveal non-antagonistic evolution consistent with ancestors residing away from a boundary. (a) Ancestor-descendant pairs plotted in evolutionary change space, where all ancestors are placed at the origin and lines indicate the direction and magnitude of change in each trait in the descendant. Pairs are colored by evolutionary outcome category as in Figure 2. Pairs for which statistical significance of evolutionary change was not reported are shown in transparent colors, as assignments to single-trait change categories (A1, A2) cannot be made. Pairs whose evolved HGT rate decreased to zero are plotted at the leftmost x-axis position; the change in log_10_(transfer rate) is undefined and these points are placed at the axis minimum for display. Inset shows the subset of single-mutation ancestor-descendant pairs on the same axes. (b) Counts of all ancestor-descendant pairs in each evolutionary outcome category. Filled bars indicate pairs with statistically supported evolutionary change; transparent bars indicate pairs without reported statistical tests, assigned to outcome categories based on direction of change alone.

Assigning pairs to these evolutionary outcome categories requires knowing whether each trait changed significantly. We therefore split the dataset into two subsets based on whether studies reported formal statistical tests of evolutionary change. For the subset with statistical tests, significant change in both traits placed a pair in a quadrant category (Q1, Q2, or Q4), while significant change in only one trait placed it in an axis category (A1 or A2). In Figure 5a, we treat the statistically supported subset (61 pairs) as our primary dataset and present the remaining subset (55 pairs) for a comparison (with reduced visual emphasis) to reflect the greater uncertainty in category assignments.

Antagonistic evolution accounts for only 59% of outcomes in the statistically supported subset (36 of 61) and 64% in the unsupported subset (35 of 55), falling far short of universal antagonism. Notably, even among antagonistic outcomes, most show weak antagonism. Large gains in one trait with minimal loss in the other are more consistent with ancestors starting deep in the interior than near the boundary (Figure 2b). The remaining outcomes that increase at least one trait are non-antagonistic (33% of supported and 31% of unsupported), a proportion irreconcilable with classical tradeoff predictions. Finally, among statistically supported pairs, changes in only one trait (A1, A2) account for 31% (19 of 61), and synergistic outcomes for less than 2% (1 of 61). There are 5 evolutionary outcomes that fall into Q3 or show no change in either trait, which we exclude from further analysis as they lack selective benefit in VGT and HGT traits. Together, these patterns are consistent with plasmid-host pairs occupying interior positions in trait space rather than residing on a boundary.

### Weak genetic constraints on plasmid transmission evolution

The evolutionary outcomes analyzed above show that a substantial proportion of plasmid-host pairs do not evolve antagonistically as expected under a tradeoff, consistent with ancestors residing in the interior of trait space. However, most of these are multi-mutation trajectories (84%, 98 of 116) and cannot tell us whether individual mutations are genetically constrained to antagonistic pleiotropy. Even interior-dwelling pairs could experience a strict genetic constraint at the mutational level, with every accessible mutation restricted to antagonistic pleiotropy. Testing this requires examining single mutants directly. A small subset of the ancestor-descendant pairs in our data set represent single mutations (Figure 5a inset, 18 pairs). Despite the limited sample size, 13 have statistical support, of which 4 show no significant change in either trait and are excluded from further analysis. Of the remaining 9, non-pleiotropic mutations dominate, accounting for 6, while antagonistic pleiotropy accounts for only 3. Thus, genetic constraints on individual mutations appear weak, with accessible mutations not restricted to antagonistic effects. We note that synergistic mutations are notably absent from this subset.

### First synergistic mutation supports HGT-driven boundary departure

To determine whether synergistic mutations are possible in plasmid transmission evolution, we returned to one of our earlier evolution experiments, whose design is well-suited to testing the HGT-driven boundary departure mechanism^21^. The original study suggested that plasmid fitness cost had decreased, implying increased VGT, and that transfer rates had qualitatively increased, but were not measured quantitatively. This made the experiment a strong candidate for documenting a synergistic outcome. The IncP-1β plasmid pB10 was introduced into a non-coevolved host, *Stenotrophomonas maltophilia*, that had been pre-adapted to laboratory conditions for 100 generations prior to plasmid introduction. The experimental protocol explicitly selected for both VGT through serial batch culture growth and HGT through regular forced conjugation cycles. We confirmed that evolved plasmids carried single mutations using long-read sequencing and measured both VGT and HGT directly for ancestral and evolved plasmids. A single amino acid mutation, V96A in *trbC*, which encodes a prepilin subunit of the conjugative pilus, was identified in 4 of 5 evolved lineages and was the only mutation in 3 of those.

This single mutation produced a synergistic outcome (Figure 6): HGT increased by over half an order of magnitude (2.52×10^−12^ to 9.89×10^−12^) and VGT increased from a fitness cost to a fitness benefit (w = 0.77 to w = 1.30). Both increases were statistically significant (HGT p = 0.015, VGT p = 0.0095, Welch t-test). This is the first documented synergistic mutation in plasmid transmission evolution, confirming that synergistic genetic changes are possible and reinforcing that antagonistic pleiotropy does not universally constrain plasmid transmission evolution. The synergistic outcome also places the ancestral plasmid-host pair in the interior of trait space. The interior position is consistent with HGT-driven boundary departure, where introduction into a non-coevolved host created a poorly adapted plasmid-host pair, thus away from the boundary before compensatory evolution can act.

**Figure 6.**
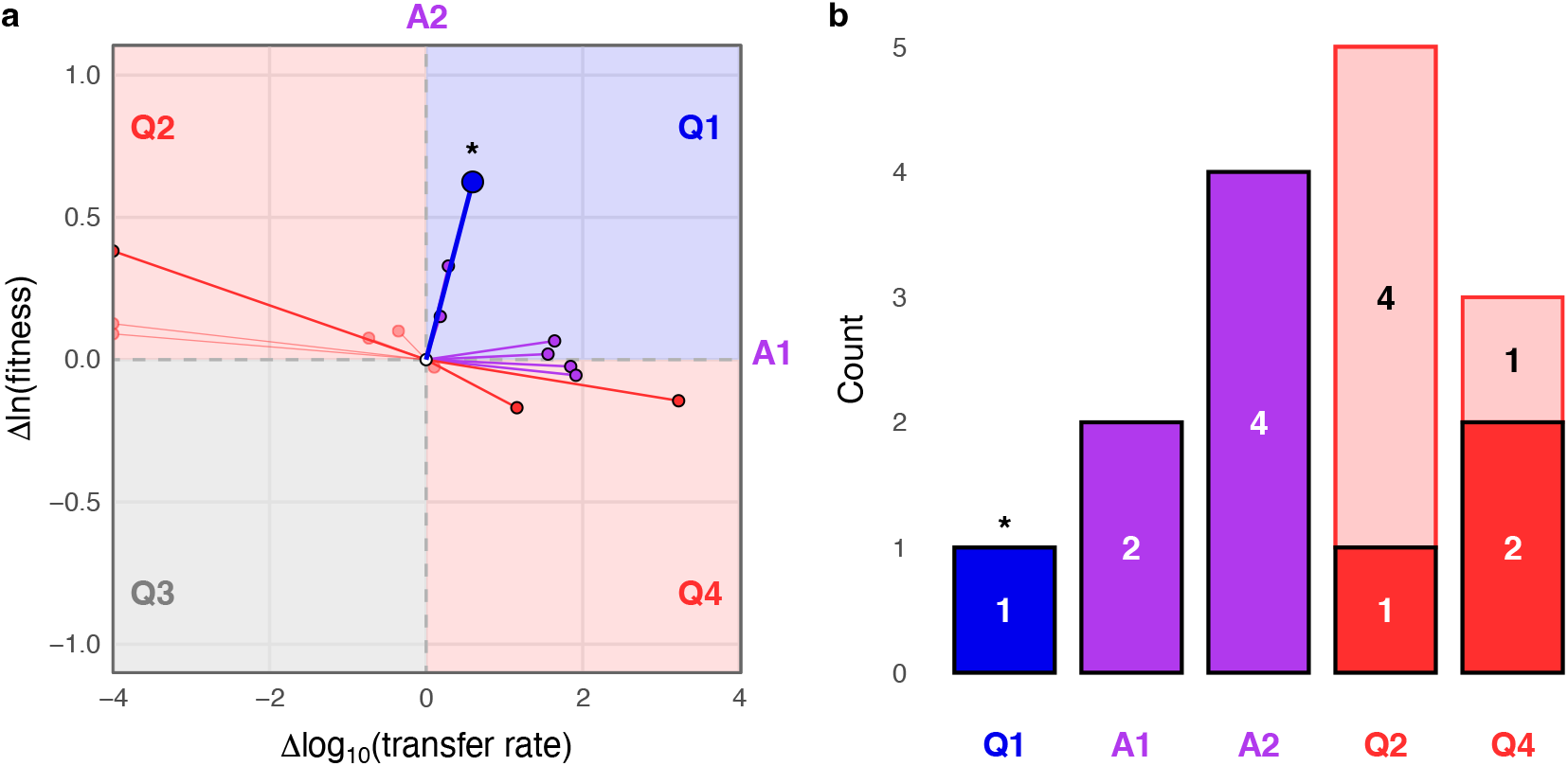
First documented synergistic mutation in plasmid transmission evolution. (a) The V96A mutation in *trbC* of the IncP-1β plasmid pB10, characterized here with new measurements of both VGT and HGT, increased both transmission traits relative to its ancestor. It is shown as a filled blue circle with an asterisk alongside the other single-mutation ancestor-descendant pairs from Figure 5a inset, with all ancestors placed at the origin and lines indicating the direction and magnitude of trait change. (b) V96A is the only synergistic outcome in the single-mutation dataset, marked with an asterisk. Counts of single-mutation ancestor-descendant pairs in each evolutionary outcome category, shown in the same format as Figure 5b.

## Discussion

Our results reveal a fundamental irony at the heart of plasmid biology: horizontal gene transfer, the defining feature of plasmid transmission, may be the very mechanism that undermines the predictive power of its own tradeoff. This is not a rejection of a VGT-HGT tradeoff, but a clarification of when the tradeoff applies. The tradeoff constrains evolution only when plasmid-host pairs are well-adapted to one another, and HGT systematically undercuts this adaptation by frequently introducing plasmids into novel hosts. The implication is that evolutionary outcomes considered inconsistent with a tradeoff are not only theoretically accessible but, as we demonstrate, empirically realized. The VGT-HGT tradeoff is therefore not a reliable predictor of plasmid evolution in general, but only following longer periods of evolution between a plasmid and its host.

The core contribution of our framework is to make explicit when a tradeoff should be expected to constrain evolution, and when it is not (Figures 1 and 2). By placing VGT and HGT in a shared trait space, the framework clarifies that a tradeoff constrains evolution only for well-adapted plasmid-host pairs residing on the boundary, while non-antagonistic outcomes arise for pairs in the interior of possible trait space. The distinction between whether a tradeoff exists and whether it constrains evolutionary outcomes has long been recognized but rarely formalized^26,36^. This gap has fueled a persistent debate over whether tradeoffs apply universally, sustained by inconsistency in what counts as evidence: tradeoffs are often assumed rather than demonstrated, and the empirical signatures used to detect them differ, with the presence of one not implying the others^37,38^. Such disagreement may not reflect genuine biological differences but variation in which signature was measured and where in trait space the sampled lineages happened to reside. Our framework reframes the problem in terms of position in trait space, replacing the ambiguous question of whether a tradeoff exists with the more tractable one of how strongly a given lineage is constrained in its evolutionary options^39,40^.

This positions our work within a broader movement toward data-driven, trait-space approaches to evolutionary tradeoffs, which characterize where organisms fall relative to a performance boundary^23,41^, and how they move toward it under selection^42^. Our framework shares this empirical grounding but differs in a crucial respect. Where existing work treats boundary residency as the endpoint that selection drives lineages toward, we make explicit the processes that move lineages off the boundary. Specifically, we identify HGT as an intrinsic boundary-departing mechanism unique to the plasmid system (Figure 3). There are dimensions we do not explicitly address that have been recently explored in other systems such as different boundary shapes^43^, soft and hard tradeoffs^44^, the role of biological innovation^45,46^, and behavior in higher-dimensional trait spaces^22,23^. These remain important open questions and our framework could be extended in the future to incorporate them.

Our cross-literature synthesis, the first to examine VGT and HGT simultaneously across diverse plasmid-host pairs, reveals patterns broadly consistent with our framework: most pairs occupy the interior of trait space rather than residing on the boundary. Interior occupancy across 16 independent studies is consistent with HGT-driven boundary departure being common rather than exceptional, though it does not on its own distinguish this mechanism from phenotypic plasticity introduced by laboratory measurement conditions, which can displace pairs from their *in situ* trait values. Disentangling these contributions, like the evidence provided by our synergistic mutation, will require measurements that track plasmid-host pairs across their coevolutionary history. The dataset also remains heavily biased toward a few replicon types and a single host species, *E. coli*, with most subgroups too sparsely sampled to support firm conclusions. Expanding the diversity of plasmids, hosts, and selective conditions will be essential for determining how reliably HGT-driven boundary departure relaxes tradeoff constraints across plasmid-host pairs.

These findings have direct implications for strategies that aim to exploit the VGT-HGT tradeoff, or similar plasmid transfer tradeoffs, to limit antibiotic resistance spread^9,12,47–49^. Such strategies assume the tradeoff reliably constrains transmission evolution, so that selection on one trait predictably comes at the expense of the other. Our framework shows this assumption is guaranteed only when plasmid-host pairs reside on the boundary, a condition HGT systematically undermines. Critically, the clinical contexts where these strategies are most needed are precisely the ones our framework predicts will fail. When resistance plasmids transfer into novel, non-coevolved hosts within patients^34,50^, the new pairs are placed in interior trait space, where evolution is no longer reliably constrained to antagonistic outcomes. Strategies premised on antagonistic constraints may therefore fail or produce unexpected outcomes in these post-transfer contexts. This does not render tradeoff-based strategies invalid. But it suggests that accounting for the coevolutionary history of the targeted pairs, and their current position in trait space, may be essential for predicting when such interventions will succeed.

More broadly, the irony we document in plasmid biology points to a general principle: the same mechanism that drives evolutionary change can be the one that relaxes its own constraints. HGT undermines the predictive power of its own tradeoff, but HGT is only one instance. The principle extends immediately to other mobile genetic elements, including phages, transposons, and integrative conjugative elements, whose horizontal movement likewise disrupts coevolutionary optimization. It extends further still to any process that displaces organisms from a tradeoff boundary, whether environmental change, phenotypic plasticity, or ecological disruption, each of which can open access to evolutionary outcomes a strict tradeoff would forbid. Where such processes are common, the classical expectation of antagonistic evolution may routinely fail, because the conditions ensuring antagonism are continually disrupted. This yields a testable prediction that reframes when we should apply the tradeoff concept: its constraints should be strongest in stable, long coevolved systems under consistent selection, and weakest in systems undergoing ecological or coevolutionary disruption. Testing this across diverse systems and trait spaces is an open frontier, with direct consequences for how confidently we will be able to use tradeoffs to predict and steer evolution.

## Acknowledgements

O.K. was supported by the NSF Postdoctoral Research Fellowships in Biology Program grant no. DBI-2305907. E.S.D. was supported by the Environmental Biology Division from the National Science Foundation (grant number 2142718). E.M.T. was supported by the Environmental Biology Division from the National Science Foundation (grant number 2142719).

---

**Extended Data Figure 1.**
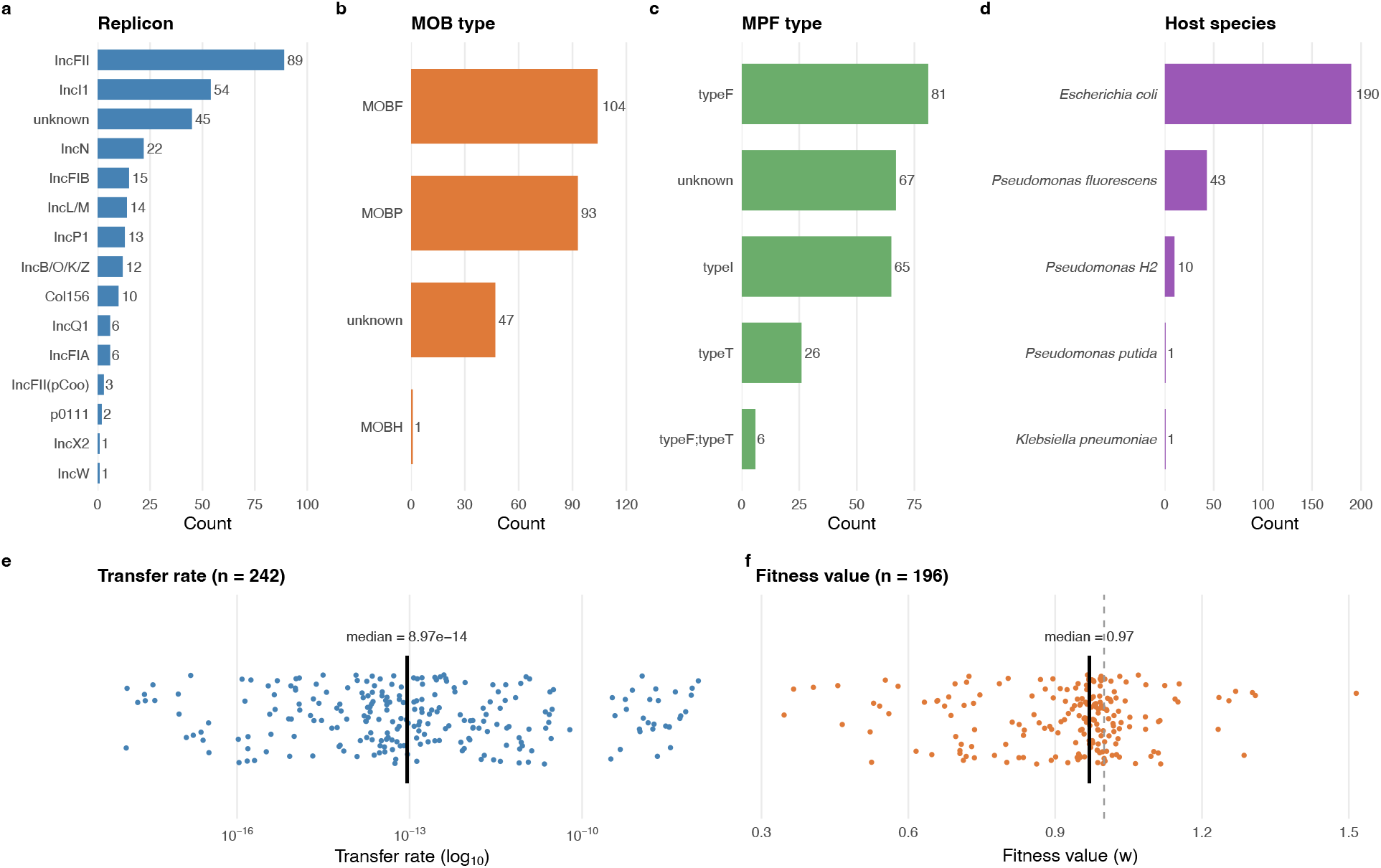
Characterization of the cross-literature dataset of plasmid-host pairs used in the VGT-HGT tradeoff analysis. (a-d) Distribution of plasmid-host pairs across biological categories: replicon type (a), MOB type (b), MPF type (c), and host species (d). Numbers indicate the count of plasmid-host pairs in each category. (e-f) Distribution of raw trait values across the dataset. Transfer rate (e, n=242) and host fitness relative to a plasmid-free host (f, n=196) are shown as dot plots with the median indicated by a solid vertical line. The dashed vertical line in panel f indicates a fitness value of w=1.0, representing no fitness effect relative to a plasmid-free host. Note that not all pairs have measurements of both traits, which is why sample sizes differ between panels e and f and why only a subset of pairs with both trait values can be plotted in trait space in Figure 4a.

**Extended Data Figure 2.**
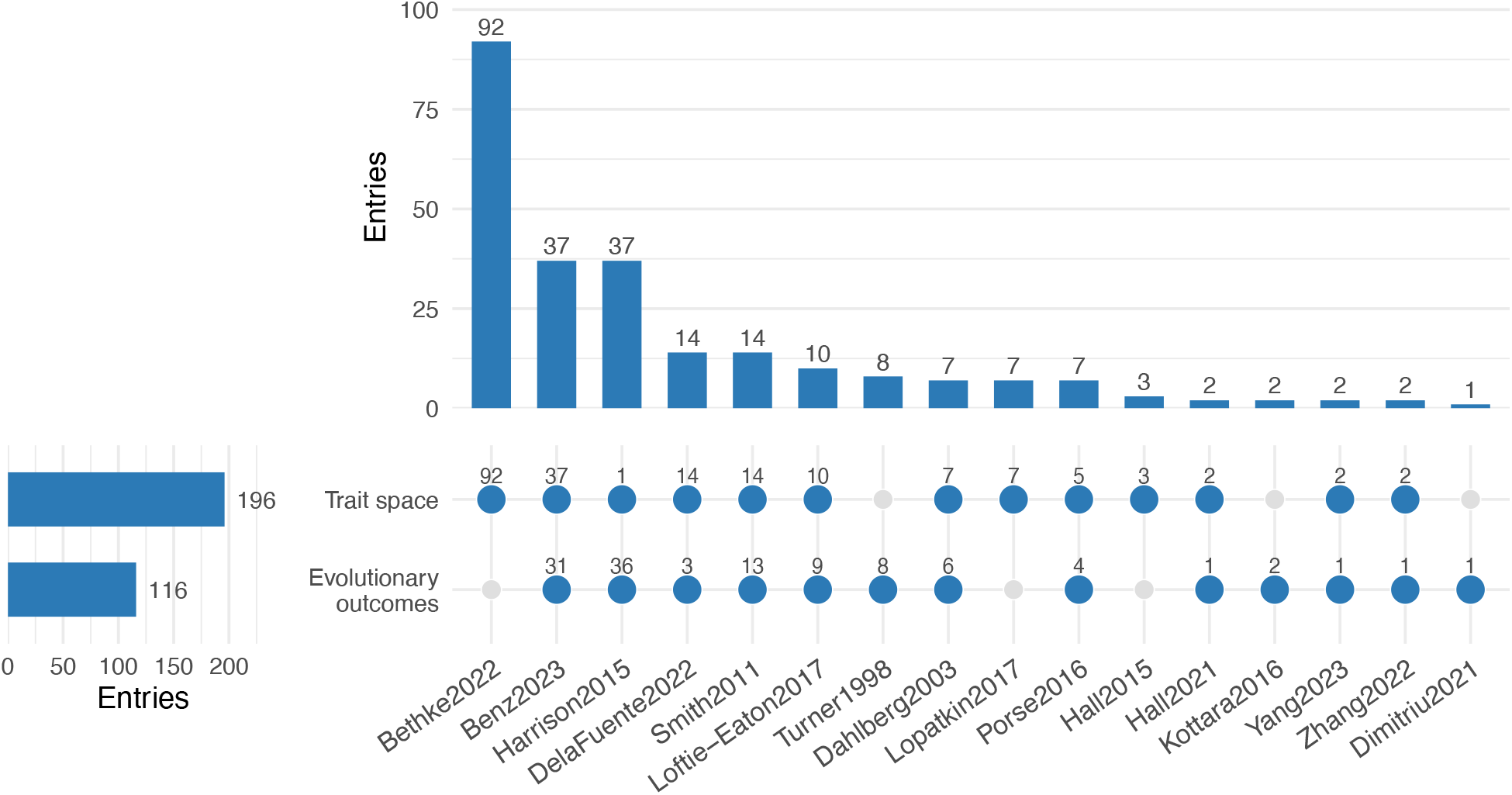
Summary of published studies contributing to the cross-literature comparison of VGT and HGT. (Top) Bar chart showing the number of plasmid-host pairs contributed by each of the 16 studies, ordered from most to fewest contributions. Studies are labeled by first author and publication year. (Bottom left) Total number of plasmid-host pairs included in each analysis. (Matrix) Filled circles indicate that a study contributes entries to the corresponding analysis; grey circles indicate no contribution. Numbers above filled circles indicate the number of entries contributed by that study. Rows correspond to the two analyses presented in the main text: trait space measurements used in Figure 4 and evolutionary outcomes used in Figures 5 and 6.

## Methods

### Literature search

We identified candidate studies through an iterative literature search using Google Scholar, Elicit, and Connected Papers, with no date or language restrictions applied. In Google Scholar, we searched for combinations of terms “plasmid” AND “fitness” AND “conjugation” and expanded these queries by substituting synonyms. For fitness, we used the terms “vertical inheritance,” “plasmid cost,” “growth rate,” and “host fitness.” For conjugation, we used “plasmid transfer,” “horizontal transfer,” and “horizontal gene transfer”. We additionally used the Elicit AI-assisted literature discovery tool, querying: “Find all primary research and review papers that explicitly examine or discuss a tradeoff or relationship between horizontal plasmid transfer (i.e., conjugation) and vertical gene transfer (e.g., host growth rate, host fitness, plasmid cost, plasmid burden).” All Elicit-returned papers were manually verified for relevance prior to screening. To capture papers linked through shared citation networks, we used Connected Papers anchored on two key studies^5,9^ and screened their associated citation graphs. Duplicates across search methods were removed prior to screening. Together these searches produced a list of 97 candidate peer-reviewed papers identified as potentially meeting inclusion criteria based on title and abstract review, of which 16 met our inclusion criteria upon full-text screening (Supplementary Table 1). This search was completed in November 2025.

### Study inclusion criteria

We included peer-reviewed studies that reported both a conjugation rate estimate and a fitness measurement for the same bacterium–plasmid pair. Conjugation rate measurements were restricted to methods that yield an estimable transfer rate parameter: the T/(D*R) method^51^, the Simonsen endpoint method^52^, the approximate Simonsen method^53^, and the Luria–Delbrück method^54^. Studies reporting only transfer frequencies were excluded, as transfer frequencies do not yield a rate estimate and are not comparable across experimental contexts^55^. Donor and recipient strains were required to be the same species.

Fitness measurements were required to be convertible to relative Malthusian fitness. We accepted two competitive reference frames. Fitness measured relative to the plasmid-free strain reflects the effect of carrying the plasmid and allows direct placement in the two-dimensional trait space used in this study, where the vertical axis represents host fitness relative to a plasmid-free cell and the horizontal axis represents plasmid transfer rate. Fitness measured relative to the plasmid-bearing ancestor instead yields the evolutionary change in fitness between ancestor and descendant (Δfitness), which captures the direction and magnitude of evolutionary change and is used for evolutionary outcome classification. Assay formats included pairwise competition experiments tracking competitor frequency over time, as well as growth rate measurements; all were converted to Malthusian fitness units (see Supplementary Table 1). Plasmids were required to be self-transmissible, encoding their own conjugation machinery. Mobilizable plasmids relying on conjugation machinery supplied in trans by a helper element were excluded, as their transfer rate reflects the helper system rather than an intrinsic property of the focal plasmid.

### Data extraction and standardization

#### Digital extraction of figures

Raw data were available for three studies; for the remaining studies, data were extracted from published figures using the web-based digitization tool Automeris. The papers and specific figures subjected to digital extraction are listed in Supplementary Table 1. Statistical significance for each evolutionary outcome was determined from information reported in the original publications using the authors’ own statistical tests, either from figure annotations or from significance statements in the text. Because individual data points and replicate-level data could not be reliably extracted across all studies, we did not rerun statistical comparisons between ancestor and descendant measurements.

#### Fitness value and transfer rate standardization

Fitness estimates were derived from either pairwise competition assays or growth rate measurements; the method used for each study is recorded in Supplementary Table 1. Although these assay types are not strictly equivalent, both yield estimates of relative Malthusian fitness. Transfer rates were used as reported in each study without further transformation, as the accepted rate-estimation methods (TDR, SIM, ASM, LDM) produce values on a common scale.

#### Replicon, MOB, and MPF typing

Plasmid replicon type, MOB class, and MPF type were assigned using COPLA^56^, a sequence-based plasmid classification tool. Where plasmid sequences were available, COPLA output was used as the primary classification. Where sequencing data were unavailable, we recorded author-reported replicon designations. MOB and MPF assignments followed the same priority using COPLA where available, otherwise inferred from author-reported incompatibility group or related sequence information when this was sufficient to make a determination. The remaining cases were recorded as NA. Plasmids carrying multiple replicons were included in each of their replicon’s analyses, but contributed a single data point to overall correlation analyses.

### Evolutionary outcome classification

Each bacterium–plasmid pair for which both a transfer rate and a fitness measurement were available before and after evolution was assigned an evolutionary outcome based on the direction of change in each trait relative to the ancestral state. Change in transfer rate (Δtransfer) and change in fitness (Δfitness) were calculated for each evolved lineage, and the sign of each delta determined quadrant placement (Figure 5): Q1 (↑transfer, ↑fitness), Q2 (↓transfer, ↑fitness), Q3 (↓transfer, ↓fitness), and Q4 (↑transfer, ↓fitness). All data entries with sufficient information for delta calculation were classified; no significance threshold was applied to quadrant assignment itself.

Only for studies in which the original authors reported statistical tests on both traits, outcomes could additionally be assigned to i) axis categories where a significant change was detected in one trait but not the other, or to a ii) no-change category where neither delta was statistically significant. When the original study did not report statistics for one or both traits, axis and no-change classification was not possible and the outcome was assigned to a quadrant based on direction of change alone. These two groups are distinguished throughout.

Q3 outcomes are included in Supplementary Table 1 but are not visualized or discussed in the main text, as they represent a small minority of classified outcomes and do not bear directly on the central question of evolutionary tradeoffs between the two traits.

### Statistical analysis

All statistical analyses were performed in R. For the meta-analysis, we assessed the relationship between plasmid transfer rate and host fitness using ordinary least squares linear regression of fitness on log_10_-transformed transfer rate, with the slope and 95% confidence interval reported. The overall Pearson correlation coefficient was calculated as a one-sided test (alternative hypothesis: negative correlation). Subgroup regressions were run independently within each replicon type, MOB class, and host species group using the same model. Subgroups with fewer than 10 data points were not evaluated. Significance of subgroup slopes was assessed at p < 0.05.

For the experimental data, transfer rates were compared between the ancestral and evolved strain using a Welch two-sample t-test on transfer rate and fitness values. Mann–Whitney U tests were run as nonparametric checks for both comparisons. Both resulted in statistical significance for the changes in VGT and HGT.

### Bacterial strains and culture conditions

All strains were derived from those used in De Gelder et al.^21^, following the same strain nomenclature, wherein P21 refers to *Stenotrophomonas maltophilia* strain P21 and “anc” denotes the ancestral host background; rifampicin- and nalidixic acid-resistant variants are denoted P21ancRif and P21ancNal, respectively. The ancestral plasmid-bearing strain consisted of the broad-host-range IncP-1β plasmid pB10, originally isolated from a wastewater treatment plant, introduced by conjugation into P21ancRif. The evolved plasmid was obtained from lineage B of the P21 evolution experiment described in De Gelder et al. Both ancestral and evolved plasmids were mated into a fresh P21ancRif background prior to all assays to ensure a uniform host background. Long-read whole-plasmid sequencing of the evolved plasmid confirmed the previous finding that it carries a single point mutation in *trbC* relative to the ancestral plasmid (Plasmidsaurus, Eugene, OR). The plasmid-free strain P21ancNal served as both the recipient in conjugation assays and the competitor in fitness assays. All strains were cultured in lysogeny broth (LB) at 30°C with shaking at 200 rpm. Tetracycline was used at 75 μg/ml for liquid selective culture and at 50 μg/ml for selective plating; nalidixic acid was used at 50 μg/ml throughout.

### Fitness assays

Pairwise competition assays were used to estimate relative fitness. All liquid culturing was carried out in 96-well microtiter plates at 30°C with continuous shaking at 200 rpm. Before each assay, strains were revived from glycerol stocks in LB overnight and subsequently transferred into fresh LB for an additional 24 hours. Plasmid-bearing and plasmid-free strains were then mixed at equal proportions in LB and allowed to compete for 24 hours. Each strain comparison was conducted in four biological replicates. Samples taken at the beginning and end of the competition period were serially diluted and deposited as spots onto LB agar containing either tetracycline (50 μg/ml) or nalidixic acid (50 μg/ml) to selectively enumerate plasmid-bearing and plasmid-free cells, respectively. Plates were incubated for 48 hours before colony counting. Relative fitness was expressed as the ratio of Malthusian parameters.

### Conjugation assays

Transfer rates were estimated using the Luria–Delbrück method^54,57^, a stochastic framework that estimates the conjugation rate from the proportion of transconjugant-free co-cultures across parallel mating assays. The LDM was implemented following all five phases of the published protocol, including verification of selective medium inhibitory concentrations, extinction probability assays, and confirmation of exponential growth during the mating window.

Strains were recovered from glycerol stocks into LB supplemented with tetracycline (75 μg/ml) for plasmid-bearing strains and into LB alone for the plasmid-free recipient, and passaged through a second overnight culture to acclimate to laboratory conditions. Cultures were then diluted and incubated for 3 hours at 30°C with shaking at 200 rpm to ensure exponential growth before mating. Donor and recipient cultures were mixed at equal volumes at a 1:1 ratio, and co-cultures were dispensed into 96 deep-well microtiter plates and incubated for 1 hour at 30°C with shaking at 200 rpm. Initial and final donor and recipient densities were estimated by dilution plating onto LB supplemented with tetracycline (50 μg/ml) or nalidixic acid (50 μg/ml), respectively. Transconjugant-selecting medium (LB supplemented with tetracycline at 100 μg/ml and nalidixic acid at 50 μg/ml) was added to all co-culture wells at the end of the mating period, and well turbidity was recorded at 72 hours. Four replicate LDM plates were run for each strain.

